# Medial prefrontal cortex population activity is plastic irrespective of learning

**DOI:** 10.1101/027102

**Authors:** Abhinav Singh, Adrien Peyrache, Mark D. Humphries

## Abstract

The prefrontal cortex is thought to learn the relationships between actions and their outcomes. But little is known about what changes to population activity in prefrontal cortex are specific to learning these relationships. Here we characterise the plasticity of population activity in the medial prefrontal cortex of male rats learning rules on a Y-maze. First, we show that the population always changes its patterns of joint activity between the periods of sleep either side of a training session on the maze, irrespective of successful rule learning during training. Next, by comparing the structure of population activity in sleep and training, we show that this population plasticity differs between learning and non-learning sessions. In learning sessions, the changes in population activity in posttraining sleep incorporate the changes to the population activity during training on the maze. In non-learning sessions, the changes in sleep and training are unrelated. Finally, we show evidence that the non-learning and learning forms of population plasticity are driven by different neuron-level changes, with the non-learning form entirely accounted for by independent changes to the excitability of individual neurons, and the learning form also including changes to firing rate couplings between neurons. Collectively, our results suggest two different forms of population plasticity in prefrontal cortex during the learning of action-outcome relationships, one a persistent change in population activity structure decoupled from overt rule-learning, the other a directional change driven by feedback during behaviour.

**Significance statement:** The prefrontal cortex is thought to represent our knowledge about what action is worth doing in which context. But we do not know how the activity of neurons in prefrontal cortex collectively changes when learning which actions are relevant. Here we show in a trial-and-error task that population activity in prefrontal cortex is persistently changing, irrespective of learning. Only during episodes of clear learning of relevant actions are the accompanying changes to population activity carried forward into sleep, suggesting a long-lasting form of neural plasticity. Our results suggest that representations of relevant actions in prefrontal cortex are acquired by reward imposing a direction onto ongoing population plasticity.

## Introduction

Among the myriad roles assigned to the medial prefrontal cortex a common thread is that it learns a model for the statistics of actions and their expected outcomes, in order to guide or monitor behaviour (Alexander and Brown, 2011; Euston et al., 2012; Holroyd and McClure, 2015; Khamassi et al., 2015; Starkweather et al., 2018; Wang et al., 2018). One way to probe this role is to use rule-switching tasks that depend on trial-and-error to uncover the statistics of each new action-outcome association. Previous work has shown that inactivating medial prefrontal cortex impairs the learning of new rules (Ragozzino et al., 1999,a; Rich and Shapiro, 2007; Floresco et al., 2008), and single pyramidal neurons change their firing times relative to ongoing theta-band oscillations only with successful rule learning (Benchenane et al., 2010). In well-trained animals, a shift in their behavioural strategy in response to a rule change is preceded by a shift in population activity in prefrontal cortex (Durstewitz et al., 2010; Karlsson et al., 2012; Powell and Redish, 2016), consistent with a change to a statistical model of the current action-outcome dependencies.

We know little though about how prefrontal cortex population activity changes during the initial learning of rules (Peyrache et al., 2009; Tavoni et al., 2017; Maggi et al., 2018). The changes to population activity could be continuous or constrained only to periods of overt learning. And these changes could be modulations of firing rates, of firing correlations, or of precise co-spiking between neurons. Knowing the continuity and form of plasticity in population activity would provide strong constraints on theories for how statistical models of the world are acquired and represented by medial prefrontal cortex.

To address these questions, here we analyse the continuity and form of population plasticity in the prefrontal cortex of rats learning rules on a Y-maze (Peyrache et al., 2009). We report that the structure of the population’s activity markedly changes between the periods of sleep either side of training on the maze. This turnover in neural activity occurs whether or not there is behavioural evidence of learning during training, and can be accounted for entirely by changes to the excitability of individual neurons, with no contribution from changes to correlations. Unique to bouts of learning is that changes to the structure of population activity in training are carried forward into the following periods of sleep. These conserved activity states are created by a combination of changes to individual neurons’ excitability and to rate, but not spike, correlations between neurons. Thus, prefrontal cortex population activity undergoes constant plasticity, but this plasticity only has a persistent direction during learning.

## Materials and Methods

### Task and electrophysiological recordings

Four Long-Evans male rats with implanted tetrodes in prelimbic cortex were trained on a Y-maze task (Figure 1A). Each recording session consisted of a 20-30 minute sleep or rest epoch (pre-training epoch), in which the rat remained undisturbed in a padded flowerpot placed on the central platform of the maze, followed by a training epoch, in which the rat performed for 20-40 minutes, and then by a second 20-30 minute sleep or rest epoch (post-training epoch). Figure 1B shows the structure of these three epochs in the ten identified learning sessions. Every trial in the training epoch started when the rat left the beginning of the departure arm and finished when the rat reached the end of one of the choice arms. Correct choice was rewarded with drops of flavoured milk. Each rat had to learn the current rule by trial-and-error, either: go to the right arm; go to the cued arm; go to the left arm; go to the uncued arm. To maintain consistent context across all sessions, the extra-maze light cues were lit in a pseudo-random sequence across trials, whether they were relevant to the rule or not.

**Figure 1.**
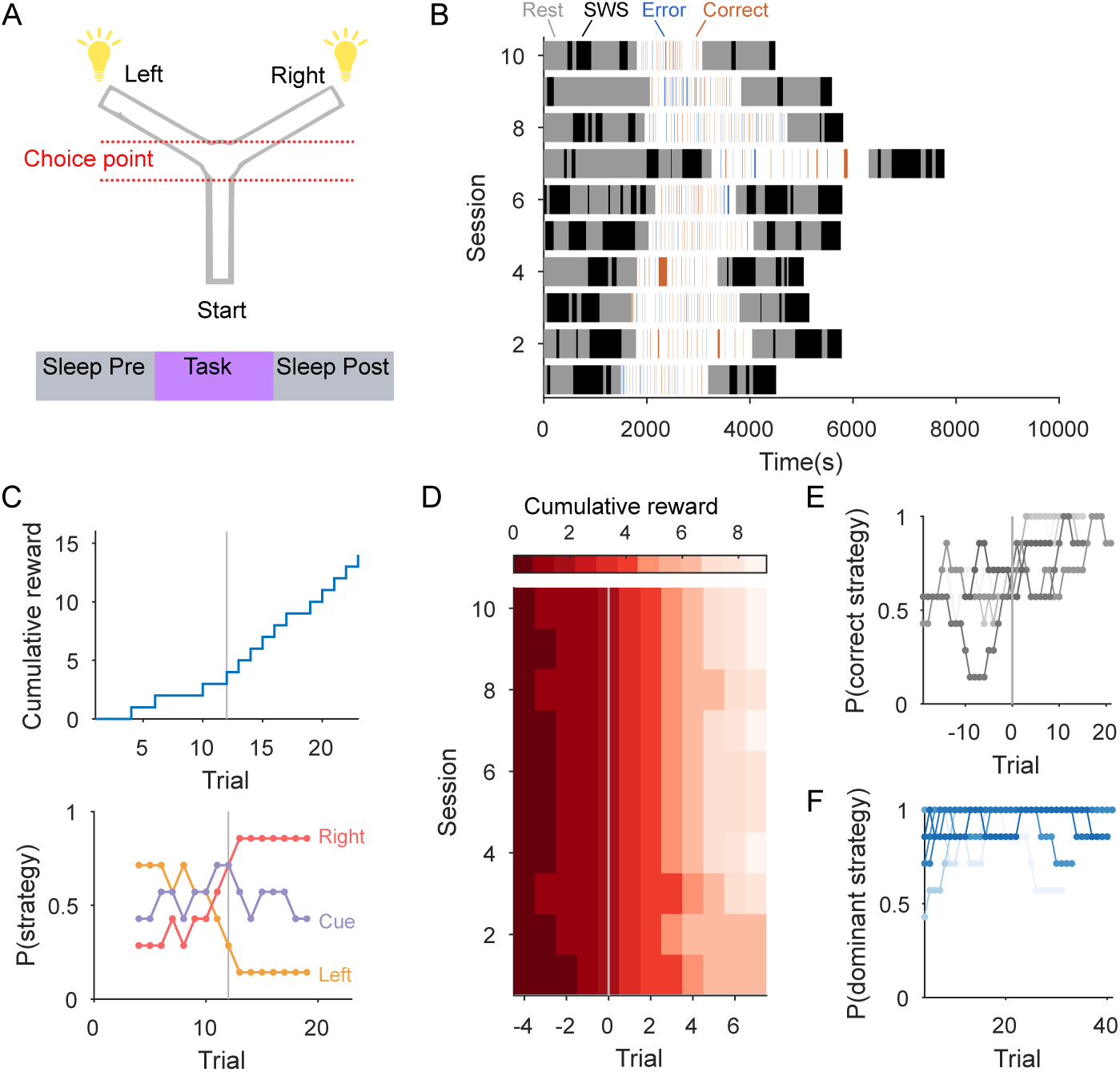
Task and behaviour. (A) Y-maze task set-up (top); each session included the epochs of pre-training sleep/rest, training trials, and post-training sleep/rest (bottom). One of four target rules for obtaining reward was enforced throughout a session: go right; go to the cued arm; go left; go to the uncued arm. No rat successfully learnt the uncued-arm rule. (B) Breakdown of each learning session into the duration of its components. The training epoch is divided into correct (red) and error (blue) trials, and inter-trial intervals (white spaces). Trial durations were typically 2-4 seconds, so are thin lines on this scale. The pre- and post-training epochs contained quiet waking and light sleep states (“Rest” period) and identified bouts of slow-wave sleep (“SWS”). (C) Internally-driven behavioural changes in an example learning session: the identified learning trial (grey line) corresponds to a step increase in accumulated reward and a corresponding shift in the dominant behavioural strategy (bottom). The target rule for this session is ‘go right’. Strategy probability is computed in a 7-trial sliding window; we plot the mid-points of the windows. (D) Peri-learning cumulative reward for all ten identified learning sessions: in each session, the learning trial (grey line) corresponds to a step increase in accumulated reward. (E) Peri-learning strategy selection for the correct behavioural strategy. Each line plots the probability of selecting the correct strategy for a learning session, computed in a 7-trial sliding window. The learning trial (grey vertical line) corresponds to the onset of the dominance of the correct behavioural strategy. (F) Strategy selection during stable behaviour. Each line plots the probability of selecting the overall dominant strategy, computed in a 7-trial sliding window. One line per session.

The data analysed here were from a total set of 50 experimental sessions taken from the study of (Peyrache et al., 2009), representing training sessions starting from naive until either the final training session, or until choice became habitual across multiple consecutive sessions (consistent selection of one arm that was not the correct arm). The four rats respectively had 13, 13, 10, and 14 sessions. From these we have used here ten learning sessions and up to 17 “stable” sessions (see below).

Tetrode recordings were spike-sorted only within each recording session for conservative identification of stable single units. In the sessions we analyse here, the populations ranged in size from 15-55 units. Spikes were recorded with a resolution of 0.1 ms. For full details on training, spike-sorting, sleep identification, and histology see (Peyrache et al., 2009).

### Session selection and strategy analysis

We primarily analyse here data from the ten learning sessions in which the previously-defined learning criteria (Peyrache et al., 2009) were met: the first trial of a block of at least three consecutively rewarded trials after which the performance until the end of the session was above 80%. In later sessions the rats reached the criterion for changing the rule: ten consecutive correct trials or one error out of 12 trials. By these criterion, each rat learnt at least two rules.

We also sought sessions in which the rats made stable choices of strategy. For each session, we computed *P*(*rule*) as the proportion of trials in which the rat’s choice of arm corresponded to each of the three rules (left, right, cued-arm). Whereas *P*(*left*) and *P*(*right*) are mutually exclusive, *P*(*cued – arm*) is not, and has an expected value of 0.5 when it is not being explicitly chosen because of the random switching of the light cue. A session was deemed to be “stable” if *P*(*rule*) was greater than some threshold *θ* for one of the rules, and the session contained at least 10 trials (this removed only two sessions from consideration). Here we tested both *θ* = 0.9 and *θ* = 0.85, giving *N* = 13 and *N* = 17 sessions respectively. These also respectively included 2 and 4 of the rule-change sessions. For the time-series in Figure 1C,E,F we estimated *P*(*rule*) in windows of 7 trials, starting from the first trial, and sliding by one trial.

### Characterising population activity as a dictionary

For a population of size *N*, we characterised the instantaneous population activity from time *t* to *t* + *δ* as an *N*-length binary vector or *word*. The *i*th element of the vector was a 1 if at least one spike was fired by the *i*th neuron in that time-bin, and 0 otherwise. Throughout we test bin sizes covering two orders of magnitude, with *δ* ranging from 1 ms to 100 ms. For a given bin size, the set of unique words that occurred in an epoch defined the dictionary of that epoch. The probability distribution for the dictionary was compiled by counting the frequency of each word’s occurrence in the epoch and normalising by the total number of time bins in that epoch.

For each session we constructed three dictionaries per bin size, and their corresponding probability distributions *P*(*Epoch*): pre-session sleep *P*(*Pre*), post-session sleep *P*(*Post*), and trials during training *P*(*Trials*). To unambiguously identify sleep periods, and for comparisons with previous reports of replay in PfC (Euston et al., 2007; Peyrache et al., 2009), we used slow-wave sleep bouts for the pre- and post-session sleep dictionaries.

We built dictionaries using the number of recorded neurons *N*, up to a maximum of 35 for computational tractability. The number of neurons used in each analysis is listed in Tables 1 and 2; where we needed to use less than the total number of recorded neurons, we ranked them according to the coefficient of variation of their firing rate between the three epochs, and choose the *N* least variable; in practice this sampled neurons from across the full range of firing rates. Only two learning sessions and six stable sessions were capped in this way.

**Table 1.**
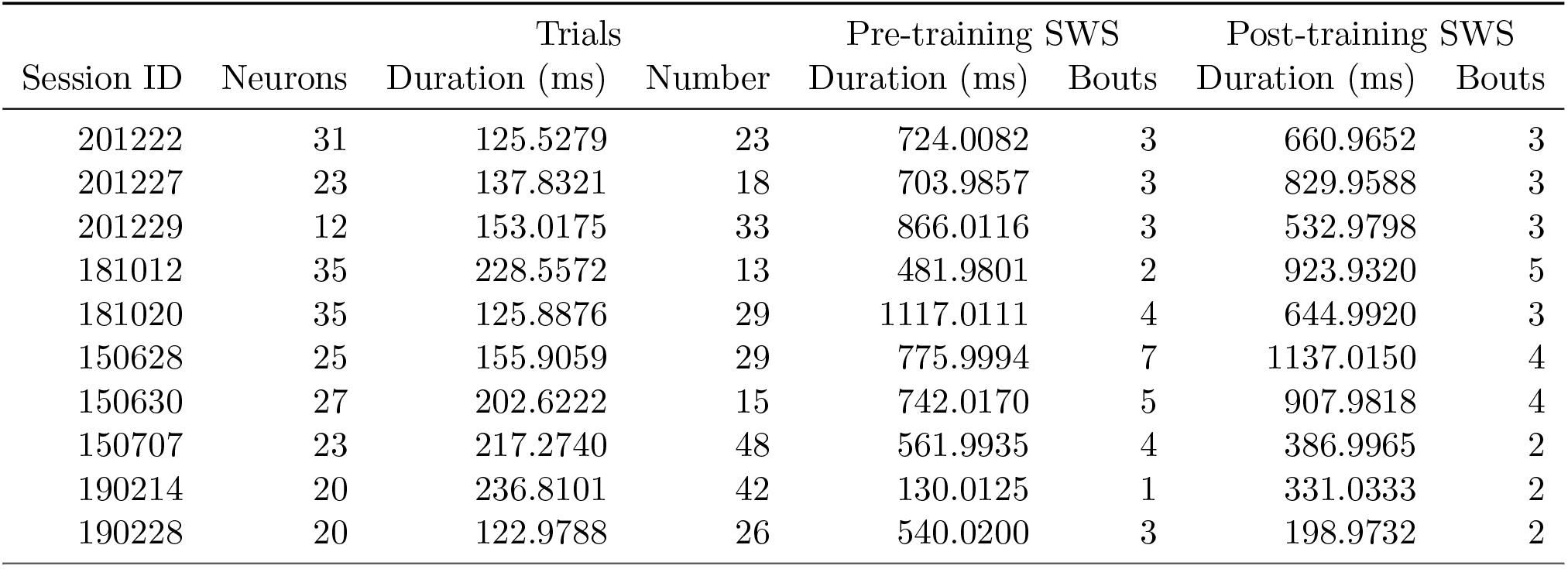
Learning session statistics. The Neurons column give the number of neurons used from each of the ten learning sessions to build the words; eight used all recorded neurons, two were capped at 35. For each epoch within a session, we give the total duration of spike-train data used to construct words, and the number of trials or sleep bouts that comprised this total duration. The number of words per epoch at a given bin size *b* can thus be calculated from this table as: Duration / *b*.

**Table 2.** Stable session statistics. Column entries as per Table 1.

| Session ID | Neurons | Trials |  | Pre-training SWS |  | Post-training SWS |  |
| --- | --- | --- | --- | --- | --- | --- | --- |
|  |  | Duration (ms) | Number | Duration (ms) | Bouts | Duration (ms) | Bouts |
| 150701 | 21 | 83.9801 | 15 | 866.0116 | 3 | 532.9798 | 3 |
| 150706 | 19 | 140.8352 | 20 | 754.0503 | 3 | 937.0184 | 3 |
| 181024 | 35 | 152.8336 | 35 | 525.9901 | 4 | 525.0188 | 2 |
| 181025 | 35 | 80.0858 | 20 | 682.0686 | 6 | 501.0109 | 4 |
| 181026 | 35 | 140.2582 | 34 | 333.0157 | 3 | 779.0119 | 5 |
| 181027 | 35 | 132.5743 | 34 | 209.9913 | 2 | 33.9931 | 2 |
| 181102 | 35 | 133.6886 | 38 | 572.9771 | 2 | 521.0275 | 7 |
| 181103 | 35 | 93.0362 | 22 | 219.9789 | 3 | 418.0025 | 4 |
| 190213 | 21 | 142.4687 | 32 | 255.9889 | 2 | 605.9930 | 1 |
| 190301 | 19 | 899.2288 | 12 | 693.0521 | 5 | 897.0089 | 5 |
| 190302 | 22 | 60.0684 | 14 | 477.0404 | 4 | 279.9953 | 1 |
| 190303 | 17 | 132.0855 | 29 | 1043.9569 | 4 | 661.0032 | 4 |
| 201228 | 19 | 217.3680 | 41 | 883.9506 | 2 | 337.9834 | 4 |
| 201230 | 21 | 171.9406 | 44 | 180.9926 | 2 | 224.9939 | 3 |
| 200102 | 22 | 195.2417 | 42 | 199.0138 | 1 | 162.0023 | 2 |
| 200103 | 29 | 289.0712 | 37 | 308.9891 | 3 | 429.9769 | 4 |
| 200105 | 12 | 223.3549 | 45 | 215.9840 | 2 | 408.0112 | 4 |

### Comparing dictionaries between epochs

We quantified the distance *D(P|Q*) between two dictionary’s probability distributions *P* and *Q* using the Hellinger distance, defined by 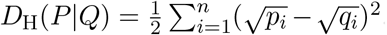. To a first approximation, this measures for each pair of probabilities (*p_i_,q*i**) the distance between their square-roots. In this form, *D_H_*(*P*|*Q*) = 0 means the distributions are identical, and *D_H_*(*P*|*Q*) = 1 means the distributions are mutually singular: all positive probabilities in *P* are zero in *Q*, and vice-versa.

To understand if a pair of pre- and post-training sleep dictionaries meaningfully differed in their structure, we compared the distance between them *D*(*Pre*|*Post*) to the predicted distance if they had an identical underlying probability distribution (in which case *D*(*Pre*|*Post*) > 0 would be solely due to finite sampling effects). We used a resampling test to estimate the predicted distance. We first created a single probability distribution *P*(*sleep*) for a session by calculating the probability of each word’s appearance in all sleep bouts across both pre and post-training sleep epochs. We then sampled *P*(*sleep*) to create new time-series of pre- and post-training sleep words, matching the number of emitted words in each epoch in the original data. By then reconstructing the dictionaries in each epoch from the resampled data, we obtained a prediction for the distance *D*(*Pre*\*|*Post*\*), where * denotes the estimate from the resampled data. Repeating the resampling 20 times gave us a distribution of expected distances assuming an identical underlying probability distribution for words. The sampling distribution’s mean and its 99% confidence interval are plotted for each session in Figure 3D,E – the intervals are too small to see on this scale.

We quantified the relative convergence of the training dictionary *X* with the dictionaries in sleep by [*D(Pre|X) – D(Post|X)]/[D(Pre|X) + D(Post|X*)]. Convergence greater than 0 indicates that the distance between the training epoch [*P(X)*] and post-training sleep [*P(Post*)] distributions was smaller than that between the training and pre-training sleep [*P(Pre*)] distributions.

### Testing hypotheses for changes in dictionary structure

To understand what drove the observed changes in the structure of population activity, we tested three hypotheses: independent changes in the excitability of neurons; changes in firing rate co-variations between neuron; and shifts in precise co-spiking between neurons. We tested these hypotheses in two steps:

1. We tested whether dictionaries constructed from independently firing neurons could account for the observed changes in the structure of population activity, with two possible outcomes:

- Yes: then we could conclude that changes in the data were due to independent changes to the excitability of the recorded neurons.
- No: this implied that the correlations between neurons were also changed.
2. To then identify the types of those correlations, we turned to dictionaries constructed from spikes jittered a little in time, and asked if they could account for the observed changes:

- No: then we would have evidence that precise co-spiking between neurons contributed to the changes in population activity structure.
- Yes: then changes to population activity did not depend on precise co-spiking, and could be accounted for by changes to co-variations in rate between neurons.

For the independent neuron dictionaries, we shuffled inter-spike intervals for each neuron independently, and then constructed words at the same range of bin sizes. As both the training and sleep epochs were broken up into chunks (of trials and slow-wave sleep bouts, respectively), we only shuffled inter-spike intervals within each chunk. This procedure kept the same inter-spike interval distribution for each neuron, but disrupted any correlation between neurons during a trial or during a sleep bout, thus testing for dictionary changes that could be accounted for solely by changes to independent neurons. We repeated the shuffling 20 times.

For any given data statistic *s_data_* for a single session, we compute the same statistic *s_shuffle_* for each shuffled data-set, and plot the difference *δ* = *s_data_* – *E*(*s_shuffle_*) using the mean *E*() over the shuffled data’s statistics. Confidence intervals at 99% for all δ were smaller than the size of the plotted symbol for *δ*, so are omitted for clarity.

For the jittered dictionaries, each spike was jittered in time by a random amount drawn from a Gaussian of mean zero and standard deviation *σ*. We tested *σ* from 2 to 50 ms. For each *σ* we constructed 20 jittered data-sets. Words were constructed from each using 5 ms bins here, both as this time-scale would capture millisecond-precise spike-timing between neurons, and because the biggest effects in the data were most consistently seen at this bin size.

We illustrated changes in the rate co-variation between neurons using the coupling between single neuron and ongoing population activity (Okun et al., 2015). Each neuron’s firing rate was the spike density function *f_i_* obtained by convolving each spike with a Gaussian of 100 ms standard deviation. Population coupling for the *i*th neuron is the Pearson’s correlation coefficient: *c_i_* = corr(*f_i_, P*_≠*i*_), where *P*_≠*i*_ is the population rate obtained by summing all firing rate functions except that belonging to the *i*th neuron.

### Relationship of location and change in word probability

To examine the spatial correlates of word occurrence, the maze was linearised, and normalised (0: start of departure arm; 1: end of the chosen goal arm). The location of every occurrence of a word during the training epoch’s trials (“trial word”) was expressed as a normalized position on the linearised maze, from which we computed the word’s median location and corresponding interquartile interval. Histograms of median word location were constructed using kernel density, with 100 equally spaced points between 0 and 1.

We tested whether the trial words closer in probability to post- than pre-training sleep were from any specific locations, which would suggest a changing representation of a key location. For each word, we computed the difference in its probability between training and pre-training sleep *δ_pre_* = |*p(pre)–p(trial)*|, and the same for post-training sleep *δ_post_* = |*p(post) – p(trial)*|, and from these computed a closeness index: (*δ_pre_ – δ_post_)/(δ_pre_* + *δ_post_*). Closeness is 0 if the word is equidistant from training to both sleep epochs, 1 if it has an identical probability between training and post-training sleep; and -1 if it has an identical probability between training and pre-training sleep.

When assessing identified maze segments, words were divided into terciles by thresholds on the closeness index at [–0.5, 0.5]; similar results were obtained if we used percentile bounds of [10, 90]%. We counted the proportion of words in each tercile whose median position fell within specified location bounds on the linearised maze. Confidence intervals on the proportions were computed using 99% Jeffrey’s intervals (Brown et al., 2001).

### Statistics

Quoted measurement values are mean *x̄* and confidence intervals for the mean [*x̄* – *t_α/2,n_SE,x̄* + *t_α/2,n_SE*], where *t_α/2,n_* is the value from the *t*-distribution at *α* = 0.05 (95% CI) or *α* = 0.01 (99% CI), and given the number *n* of data-points used to obtain *x̄*. For testing the changes in convergence, we used the Wilcoxon signed-rank test for a difference from zero; for differences in population-coupling correlations, we used the Wilcoxon signed-rank paired-sample test. Throughout, we have *n* = 10 learning sessions and *n* = 17 stable sessions.

### Data and code availability

The spike-train and behavioural data that support the findings of this study are available at CRCNS.org (DOI: 10.6080/K0KH0KH5) (Peyrache et al., 2018). The sessions meeting our learning and stable criteria are listed in Tables 1 and 2.

Code to reproduce the main results of the paper is available at: https://github.com/mdhumphries/PfCDictionary.

## Results

### Signatures of rule-learning on the Y-maze

Rats with implanted tetrodes in the prelimbic cortex learnt one of four rules on a Y-maze: go right, go to the randomly-cued arm, go left, or go to the uncued arm (Figure 1A). Rules were changed in this sequence, unsignalled, after the rat did 10 correct trials in a row, or 11 correct trials out of 12. Each rat learnt at least two of the rules, starting from a naive state. Each training session was a single day containing 3 epochs totalling typically 1.5 hours: pre-training sleep/rest, behavioural training on the task, and posttraining sleep/rest (Figure 1B). Here we consider bouts of slow-wave sleep throughout, to unambiguously identify periods of sleep. Tetrode recordings were spike-sorted within each session, giving populations of single neuron recordings ranging between 12 and 55 per session (see Tables1 and 2 for details of each session and each epoch within a session).

In order to test for the effects of learning on the structure of joint population activity, we need to compare sessions of learning with those containing no apparent learning as defined by the rats’ behaviour. In the original study containing this data-set, Peyrache et al. (2009) identified 10 learning sessions as those in which three consecutive correct trials were followed by at least 80% correct performance to the end of the session; the first trial of the initial three was considered the learning trial. By this criterion, the learning trial occurs before the mid-point of the session (mean 45%; range 28-55%). We first check this criterion corresponds to clear learning: Figure 1C,D shows that each of the ten sessions has an abrupt step change in reward accumulation around the identified learning trial corresponding with a switch to a consistent, correct strategy within that session (Figure 1E).

We further identify a set of 17 sessions with a stable behavioural strategy throughout, defined as a session with the same strategy choice (left, right, cue) on more than 85% of trials (Figure 1F). This set includes four sessions in which the rule changed. Setting this criterion to a more conservative 90% reduces the number of sessions to 13 (including two rule change sessions), but does not alter the results of any analysis; we thus show the 85% criterion results throughout.

### Constant plasticity of population activity between sleep epochs

We want to describe the joint population activity over all *N* simultaneously-recorded neurons with minimal assumptions, so that we can track changes in population activity however they manifest. Dividing time into bins small enough that each neuron either spikes (‘1’) or doesn’t (‘0’) gives us the instantaneous state of the population as the N-element binary vector or *word* in that bin (Figure 2). The dictionary of words appearing in an epoch and their probability distribution together describe the region of joint activity space in which the population is constrained. Comparing dictionaries and their probabilities between epochs will thus reveal if and how learning changes this region of joint activity.

**Figure 2.**
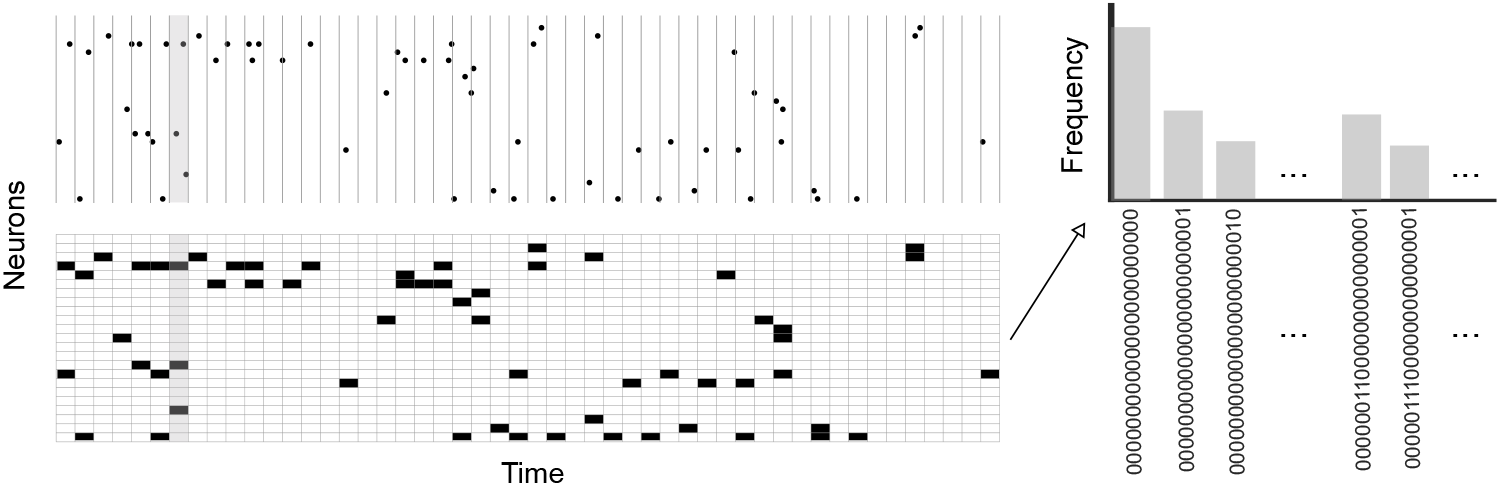
A neural dictionary of population activity in prefrontal cortex. A snapshot of population activity from *N* = 23 neurons during 500 ms of pre-training sleep, and below is the corresponding binary word structure (black: 1; white: 0) for bins of 10 ms. One bin of the population activity and its corresponding binary word is highlighted in grey. Right: The set of binary words and the frequency of their occurrence over the whole pre-training sleep epoch defines a dictionary of population activity.

If learning during training correlated with changes to the underlying neural circuit in prefrontal cortex then we might reasonably expect population activity in post-training sleep to also be affected by these changes, and so differ from activity in pre-training sleep. We thus compare the dictionaries in pre- and post-training sleep for the learning sessions, and then check if any detected changes also appear during sessions of stable behaviour.

A first check is simply if the dictionary content changed during learning and not stable behaviour. We find that the words common to both sleep epochs (Figure 3A) account for almost all of each epoch’s activity (Figure 3B) at bin sizes up to 20 ms. Consequently, there are no differences between learning and stable behaviour in the overlap of dictionary contents between sleep epochs (Figure 3A) or in the proportion of activity accounted for by words common to both sleep epochs (Figure 3B). We could thus rule out that learning changes the dictionary content between sleep epochs compared to stable behaviour. Any learning-specific change ought then be found in the structure of the population activity.

**Figure 3.**
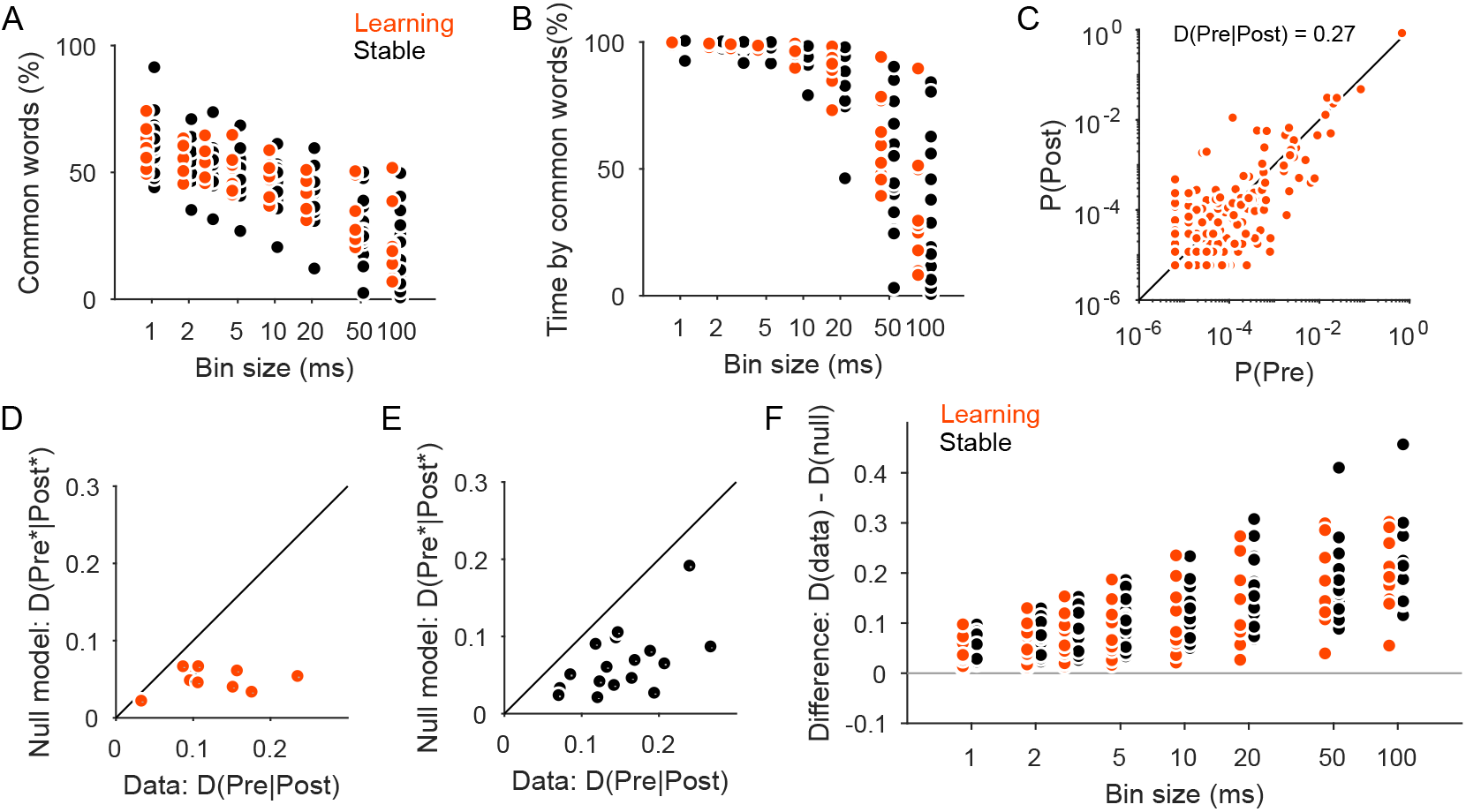
Distributions of word probabilities change between pre- and post-training sleep. (A) Proportion of words in the pre-training sleep dictionary that are also in the post-training sleep dictionary, per session. (B) Proportion of the pre-training sleep epoch’s activity that is accounted for by words in common with post-training sleep, per session. (C) The joint distribution of the probability of every word occurring in pre-training sleep (distribution P(Pre)) and post-training sleep (distribution *P*(*Post*)), for one learning session. D(Pre|Post): the distance between the two probability distributions for words. (D) Distance between the word probability distributions for pre- and post-training sleep (x-axis) against the expected distance if the sleep activity was drawn from the same distribution in both epochs (y-axis). One symbol per learning session; we plot the mean and 99% confidence interval (too small to see) of the expected distance *D(Pre*\*|*Post*\*). Words constructed using 5 ms bins. (E) Same as (D), for stable sessions. (F) Bin-size dependence of changes in the dictionary between sleep epochs. Difference between the data and mean null model distance are plotted for each session, at each bin-size used to construct words.

We capture this structure by the respective distributions *P(Pre)* and *P(Post)* for the probability of each word appearing in pre- or post-training sleep. Changes to the detailed structure of the pre- and post-training sleep dictionaries are then quantified by the distance between these probability distributions (Figure 3C). These distances will vary according to both the number of neurons *N* and the duration of each epoch. So interpreting them requires a null model for the distances expected if *P*(*Pre*) and *P*(*Post*) have the same underlying distribution *P*(*Sleep*), which we approximate using a resampling test (see Methods). In this null model any differences between *P*(*Pre*) and *P*(*Post*) are due to the finite sampling of *P*(*Sleep*) forced by the limited duration of each epoch.

In learning sessions the distance between pre- and post-training sleep probability distributions always exceeds the upper limit of the null model’s prediction (Figure 3D). This was true at every bin size (Figure 3F), even at small bin sizes where the dictionaries were nearly identical between the sleep epochs (Figure 3A). Thus, the probability distributions of words consistently differ between pre and post-training sleep epochs in learning sessions.

However, Figure 3E-F shows this consistent difference is also true for the sessions with stable behaviour. There is quantitative agreement too as the gap between the data and predicted distances has the same distribution for both learning and stable behaviour (Figure 3F). We conclude that the probabilities of words do systematically change between sleep epochs either side of training, but do so whether there is overt learning or not.

### Learning systematically updates the dictionary

This leaves open the question of whether changes in population activity between sleep epochs are a consequence of changes during training. If the population changes between sleep epochs are unrelated to population activity in training, then the probability distribution of words in training will be equidistant on average from that in pre- and post-training sleep. Alternatively, changes to population activity during training may carry forward into post-session sleep, possibly as a consequence of neural plasticity during the trials changing the region of joint activity space in which the population is constrained. A prediction of this neural-plasticity model is that the directional change would thus occur predominantly during learning sessions, so that only in these sessions is the distribution of word probabilities in training closer to that in post-training sleep than in pre-training sleep.

Unpicking the relationship between the sleep changes and training requires that the dictionary in training also appears in the sleep epochs; otherwise changes to word probabilities during training could not be tracked in sleep. We find that the structure of population activity in training is highly conserved in the sleep epochs (Figure 4A), both in that the majority of words appearing in trials also appear in the sleep epochs, and that these common words account for almost all of the total duration of the trials. This conservation of the training epoch population structure in sleep allows us to test the prediction of a learning-driven directional change in population structure (Figure 4B).

**Figure 4.**
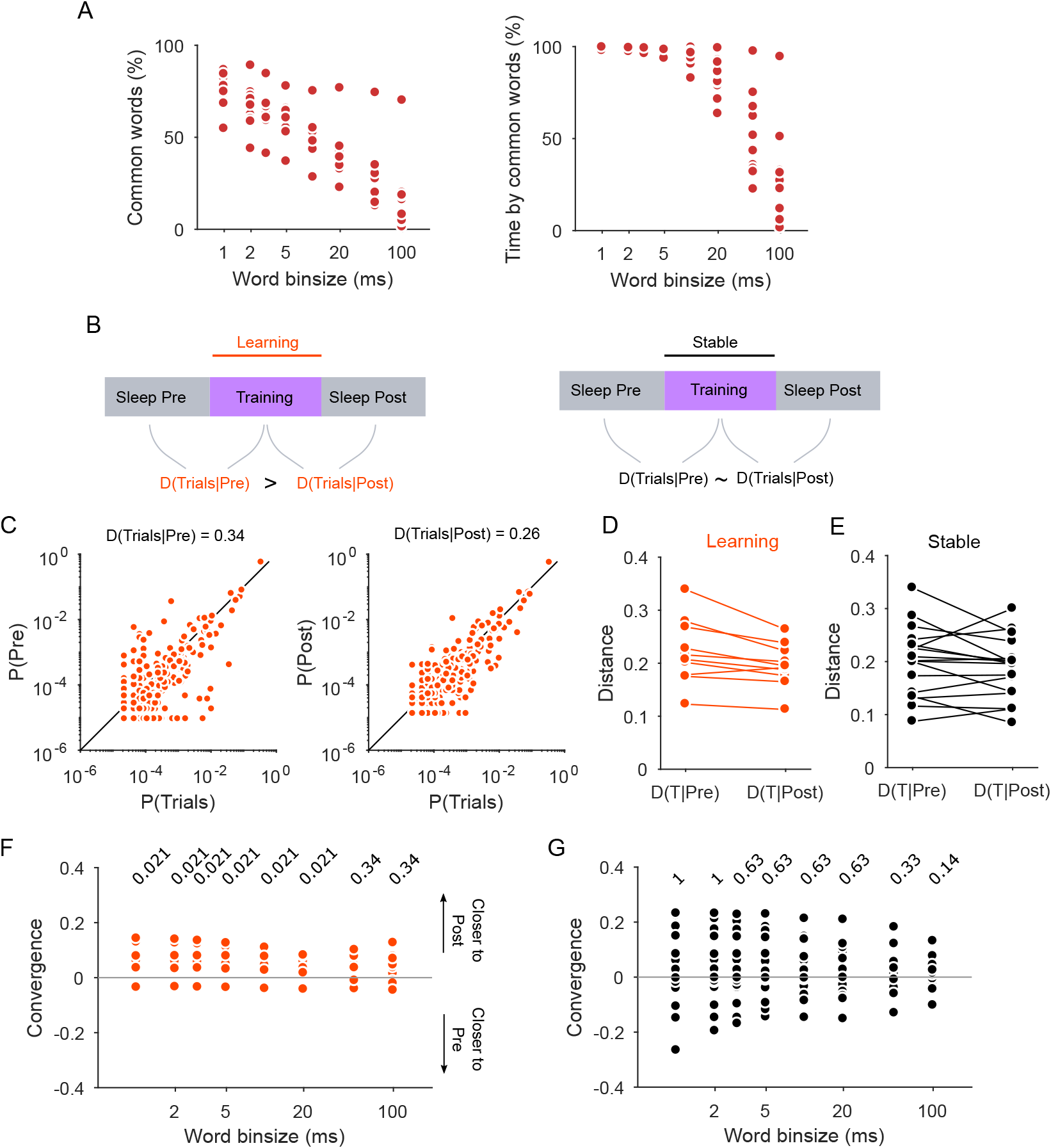
Distributions of word probabilities converge only during learning. (A) For the training epochs, the proportion of the epoch’s dictionary (left) and duration (right) accounted for by words in common with both sleep epochs. One symbol per learning session. (B) Schematic of comparisons between epochs, and summary of main results. (C) Examples for one learning session of the joint probability distributions for each word in trials and pre-training sleep (left), and trials and post-training sleep (right), using 5 ms bins. *D(Trials|X)*: the distance between the two probability distributions for words. (D) Distances for all learning sessions, for words constructed using 5 ms bins. T: Trials. (E) As for (D), for stable sessions. (F) Bin-size dependence of the relative convergence between the word distributions in trials and in sleep. Each distance was computed using only the dictionary of words appearing in the trials. Numbers are P-values from two-sided signtests for each median differing from zero. (G) As for (F), for stable sessions.

To do so, we take the dictionary of words that appear during training, and compute the distance between its probability distribution and the probability distribution of that dictionary in pre-training sleep (D(Prel Learn)), and between training and post-training sleep (*D(Post|Learn)*) (Figure 4C). The prediction of the directional change model is then *D(Pre|Learn)* > *D(Post|Learn)*. This is exactly what we found: *D(Pre|Learn*) is consistently larger than *D(Post|Learn*) at small bin sizes, as illustrated in Figure 4D for 5 ms bins.

If these directional changes are uniquely wrought by learning, then it follows that we should not see any systematic change to the dictionary in the stable behaviour sessions (Figure 4B). To test this prediction, we similarly compute the distances *D(Pre|Stable*) and *D(Post|Stable*) using the dictionary of words from the training epoch, and test if *D(Pre|Stable) ≈ D(Post|Stable*). Again, this is exactly what we found: *D(Pre|Stable*) was not consistently different from *D(Post|Stable*) at any bin size, as illustrated in Figure 4E for 5 ms bins.

It is also useful to consider not just which sleep distribution of words is closer to the training distribution, but how much closer. We express this as a convergence ratio *C* = [*D(Pre|X)* – *D(Post|X)]/[D(Pre|X*) + *D(Post|X*)], given the training distribution *X* = *{Learn, Stable}* in each session. So computed *C* falls in the range [–1,1] with a value greater than zero meaning that the training probability distribution is closer to the distribution in post-training sleep than the distribution in pre-training sleep. Figure 4G shows that for learning sessions the word distribution in training is closer to the posttraining than the pre-training sleep distribution across an order of magnitude of bin sizes. For stable sessions the absence of relative convergence is consistent across two orders of magnitude of bin size (Figure 4G). Both qualitatively and quantitatively, the structure of prefrontal cortex population activity shows a relative convergence between training and post-training sleep that is unique to learning.

### Changes to neuron excitability account for changes between sleep epochs

What then is the main driver of the observed changes in the structure of population activity? These could arise from changes to the excitability of independent neurons, to covariations in rate over tens to hundreds of milliseconds, or to the millisecond-scale precise timing of co-incident spiking between neurons. We first examine the drivers of the changes between sleep epochs we saw in Figure 3.

Individual sessions showed a rich spread of changes to neuron excitability between the sleep epochs (Figure 5A). We thus begin isolating the contribution of these three factors by seeing how much of the change in population structure between sleep epochs can be accounted for by independent changes to neuron excitability. Shuffling inter-spike intervals within each epoch gives us null model dictionaries for independent neurons by removing both rate and spike correlations between them, but retaining their excitability (at least, as captured by their inter-spike interval distribution).

**Figure 5.**
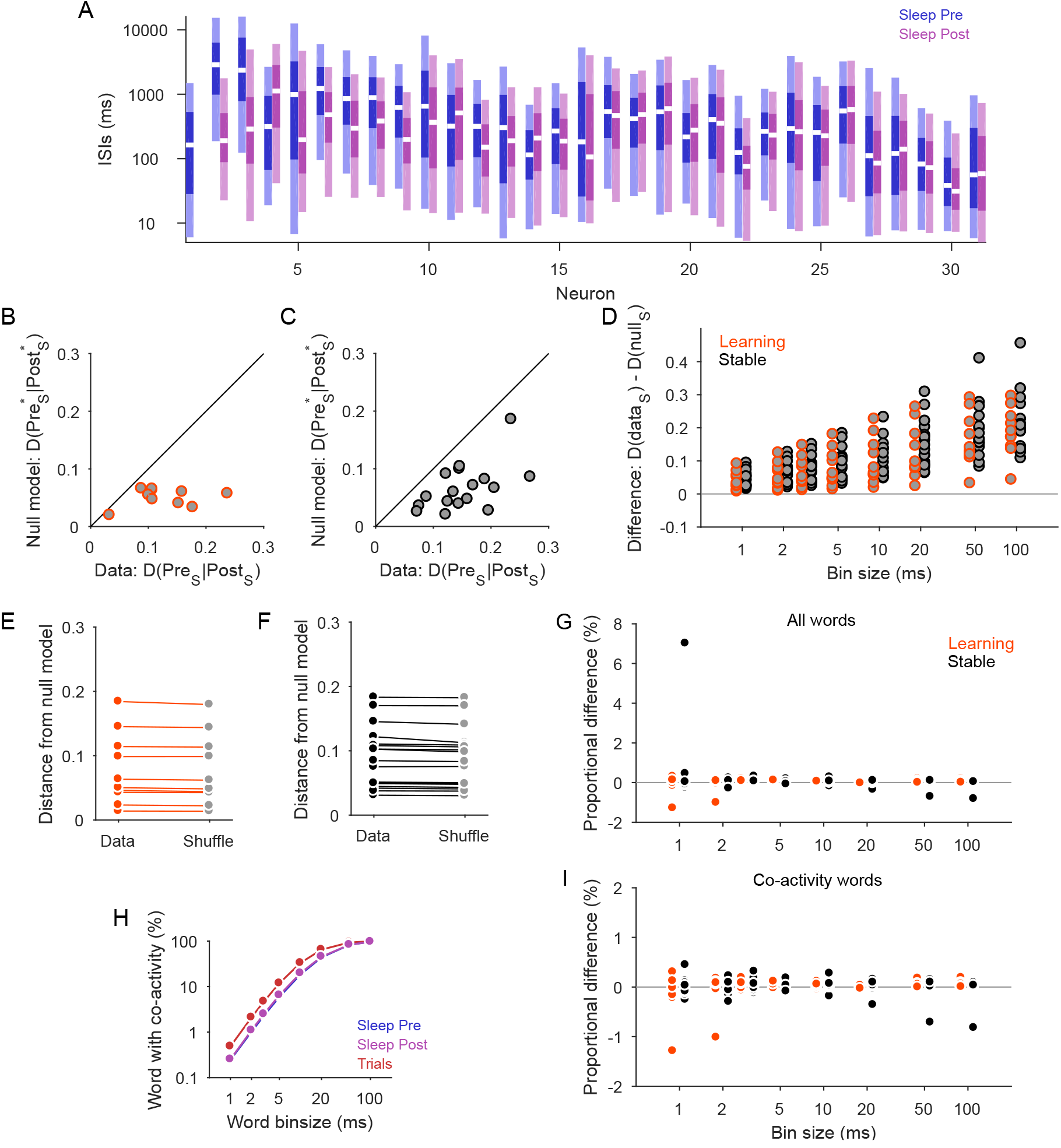
Changes between sleep epochs are accounted for by independently changing neurons. (A) Example excitability changes between sleep epochs, for one learning session. Each pair of bars plot the distributions of a neuron’s inter-spike intervals in the pre- and post-training sleep epochs, each bar showing the median (white line), interquartile range (dark shading) and 95% interval (light shading). Neurons are ranked by the difference in their median interval between sleep epochs. We use a log-scale on the y-axis: some neurons shift their distribution over orders of magnitude between sleep epochs. The first neuron was silent in the post-training sleep epoch. (B) Distances between sleep epochs for dictionaries of independent neurons (x-axis), and their expected distances from a null model of the same dictionary in both epochs (y-axis). Independent neuron dictionaries are constructed by shuffling inter-spike intervals within trials or sleep bouts. One symbol per learning session; we plot the mean and 99% confidence interval (too small to see) of the expected distance *D(Pre\** |*Post\**). Words constructed using 5 ms bins. S: shuffled data. (C) As for (B), for stable sessions. (D) Independent neuron dictionaries are consistently different between sleep epochs at all bin sizes – compare to results for the data dictionaries in Figure 3F. Each symbol is a mean over 20 shuffled data-sets. (E) Departure from the expected distance between sleep epochs for each learning session (Data), and the corresponding predicted departure by independent neurons (Shuffle; mean over 20 shuffled data-sets). Words constructed using 5 ms bins. (F) As for (D), for stable sessions. (G) Difference between the recorded and shuffled data, as a proportion of the data’s departure from the expected distance between sleep epochs. Almost all differences are less then 0.1% of the difference between data and the null model. One symbol per session. (H) The proportion of words in the dictionary with two or more active neurons, over all learning sessions. (I) As for panel (G), using dictionaries that contained only words with co-activity. At all bin sizes, there is no systematic difference between recorded and shuffled data.

When we analyse the changes between sleep epochs for independent neuron dictionaries, the strong similarity with the results from the data dictionaries is compelling. We illustrate this in Figure 5B-D, by repeating the analyses in Figure 3D-F but now using the independent neuron dictionaries – and see the results are essentially the same. The departure from the null model of a single probability distribution in sleep is almost identical between the data and independent neuron dictionaries, illustrated in Figure 5E-F for 5 ms bins. And while the data dictionaries tend to depart further from the null model, this excess is negligible, being on the order of 0.1% of the total departure from the null model (Figure 5G).

A potential confound in searching for the effects of correlation here are that words coding for two or more active neurons are infrequent at small bin sizes, comprising less then 10% of words at small bins sizes (Figure 5H). As a consequence, any differences between the independent neuron and data dictionaries that depend on correlations between neurons in the data could be obscured. To check for this, we repeat the same analyses of the changes between sleep for both the data and independent neuron dictionaries when they are restricted to include only co-activity words. As Figure 5I shows, this did not uncover any hidden contribution of correlation between neurons in the data; indeed, for coactivity words alone, the difference between the data and the independent model is about zero. Thus, the changes in word probabilities between pre- and post-training sleep can be almost entirely accounted for by independent changes to the excitability of individual neurons (Figure 5A).

### Learning-driven changes to the dictionary include rate co-variations

Can independent changes to individual neuron excitability also account for the relative convergence of dictionaries in learning? Repeating the comparisons of training and sleep epoch activity using the independent neuron dictionaries, we observe the same learning-specific convergence of the training and post-training sleep dictionaries, illustrated in Figure 6A for 5 ms bins (compare Figure 4D-E). Figure 6B shows that the difference in convergence score between the data and independent neuron dictionaries is close to zero at most bin sizes. This suggests that the changes in population activity during the trials that are carried forward to the post-training sleep can also be accounted for by the changing excitability of individual neurons.

**Figure 6.**
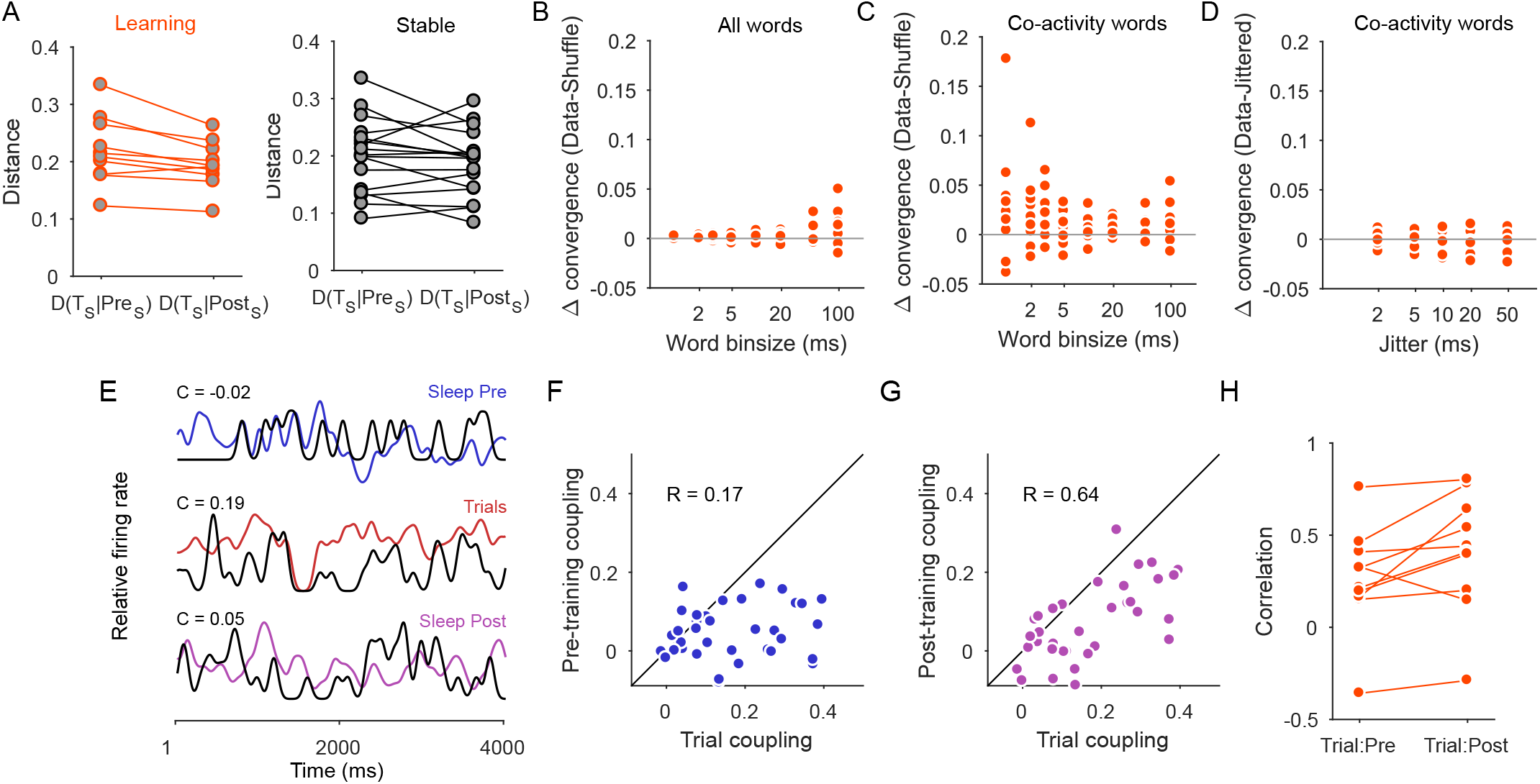
Convergence of dictionaries during learning is partly driven by changes in rate co-variation, but not spike-timing. (A) Distances between sleep and trial distributions for all learning (left) and stable (right) sessions, in an example shuffled data-set. Words constructed using 5 ms bins. *D(T_s_|X_S_)*: the distance between the trial probability distribution and the probability distribution of sleep epoch *X* in the shuffled data. (B) Difference between the recorded and shuffled data convergence between trial and post-training sleep epochs, in learning sessions. (C) As for panel B, using distributions containing only words with co-activity. (D) As for panel C, comparing co-activity word distributions from recorded and jittered data, to test for the contribution of precise spike timing. Spike data were jittered at a range of standard deviations (x-axis), and words constructed using 5 ms bins. (E) Snapshots of a single neuron’s firing rate (black) in comparison to the simultaneous population firing rate (colour) in each epoch. C: population-coupling in each epoch. (F) Joint distribution of the population coupling for each neuron in the training and pre-training sleep epochs of one learning session. R: Pearson’s correlation coefficient between the two distributions of population coupling. (G) Same as (F), for the training and post-training sleep epochs in the same session. (H) Correlations between population coupling in training and sleep epochs for all learning sessions. Population-coupling is more correlated between training and post-training sleep (signed-rank test *P* = 0.02, *rank* = 5).

To check this conclusion, we again account for the relative infrequency of co-activity words at small bin sizes by recomputing the distances between sleep and training epochs using dictionaries of only co-activity words. Now we find that, unlike the changes between sleep epochs, the relative convergence between training and post-training sleep for the data dictionaries is greater than for the independent neuron dictionaries (Figure 6C). We conclude that changes to the correlations between neurons during the trials of learning sessions are also detectably carried forward to post-training sleep.

These correlations could take the form of co-variations in rate, or precise co-incident spikes on millisecond time-scales. To test for precise co-spiking, we construct new null model dictionaries: we jitter the timing of each spike, and then build dictionaries using 5 ms bins to capture spike alignment. If precise co-spiking is contributing to the correlations between neurons, then relative convergence should be smaller for these jittered dictionaries than the data dictionaries. As Figure 6D shows, this is not what we found: across a range of time-scales for jittering the spikes, the difference in relative convergence between the data and jittered dictionaries was about zero. The changed correlations between neurons are then rate co-variations, not precise co-spiking.

Figure 6E-H gives some intuition for these changes in rate co-variation. We measure the coupling of each neuron’s firing to the ongoing population activity (Figure 6E) as an approximation of each neuron’s rate covariation (as population-coupling is fixed to a particular time-scale, so it can only represent part of the co-variation structure captured by the dictionaries of words). The distribution of population coupling across the neurons varied between epochs (Figure 6F-G), signalling changes to the co-variations in rate between neurons. Consistent with changes to rate co-variations, the distribution of coupling tended to be more similar between training and post-training sleep than between training and pre-training sleep (Figure 6H).

### Locations of dictionary sampling during learning

The changes to population activity in training carried forward to post-training sleep may correspond to learning specific elements of the task. We check for words linked to task elements by first plotting where each word in the training dictionary occurs on the maze during trials. Words cluster at three maze segments, as illustrated in Figure 7A for 3 ms bins: immediately before the choice area, at its centre, and at the end of the chosen arm. This clustering is consistent across all bin sizes (Figure 7B).

**Figure 7.**
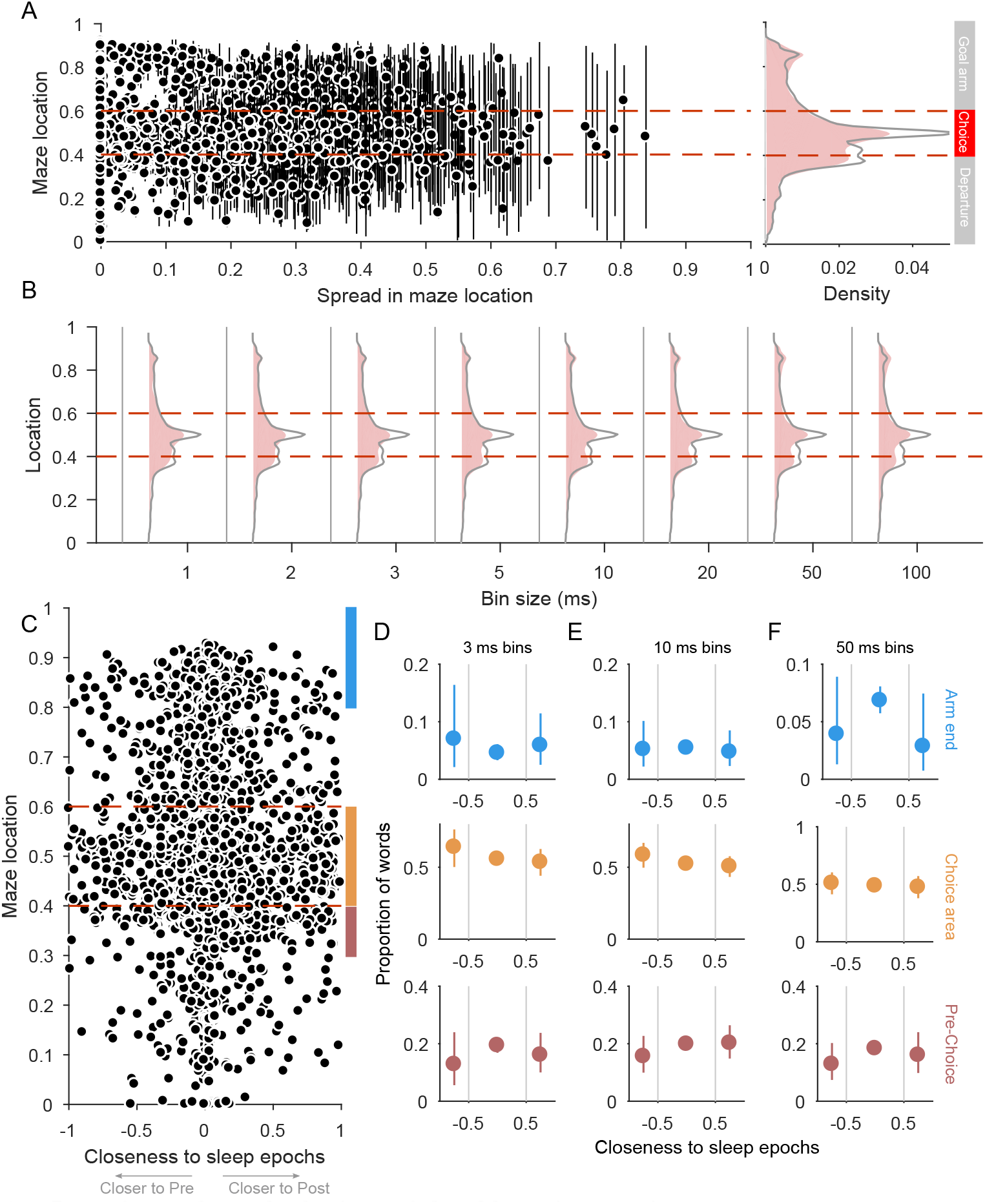
Locations of words during trials of learning sessions. (A) Scatter of the spread in location against median location for every word in the training epoch dictionaries of the learning sessions, constructed using 3 ms bins. Spread in location is the inter-quartile interval, which we also plot as vertical lines. On the right we plot the density of median locations for the data (red area plot) and independent neuron (grey line) dictionaries. (B) Density of median locations across all bin sizes, for data (red area plot) and independent neuron (grey line) dictionaries. (C) For each word in the training epoch dictionaries, we plot its median location against the closeness between its training epoch and sleep epoch probability. Closeness is in the range [-1,1], where -1 indicates identical probability between training and pre-training sleep, and 1 indicates identical probability between training and post-training sleep. Coloured bars indicate the regions of the maze analysed in panels D-F. (D) Distributions of word closeness to sleep in specific maze segments, for 3 ms bins. All words with median locations within the specified maze segment are divided into terciles of closeness by thresholds of -0.5 and 0.5 (vertical grey lines). Symbols plot proportions of words falling in each tercile, and error bars plot 99% confidence intervals on those proportions. Blue: arm end; orange: choice point; red: pre-choice segment. (E) As panel D, for 10 ms bins. (F) As panel D, for 50 ms bins.

Repeating this location analysis using the dictionaries of independent words gives the same three clusters (grey lines in Figure 7A-B). This suggests that the clustering of words at particular locations can be largely attributed to the amount of time the animals spent at those locations. The only departures are that the choice region is slightly underrepresented in the data dictionaries, and the arm-end slightly over-represented. These departures are potentially interesting, as they correspond to key points in the task: the area of the maze at which the goal arm has to be chosen, and the arrival at the goal arm’s reward port.

We thus check if words in these three segments are more likely to have their probabilities in training carried forward to post-training sleep. Figure 7C shows that when we plot the closeness of each word’s probability in training and sleep, we obtain a roughly symmetrical distribution of locations for words closer to pre-training and post-training sleep. At the three maze segments, we indeed find that a word’s probability in training is equally likely to be closer to pre-training sleep, equidistant from both sleep epochs, or closer to posttraining sleep (Figure 7D-F). We obtain the same results if we use just co-activity words, or if we divide the closeness distribution into pre/equidistant/post by percentiles rather than the fixed ranges we use in Figure 7D-F (data not shown). There is, then, no evidence in this analysis that words representing specific maze locations, and putatively key task elements, have their changes in training carried forward to post-training sleep. Rather, changes to the structure of population activity during learning are distributed over the entire maze.

### Independent neurons capture the majority of structure in prefrontal cortex population activity

The above analyses have shown that independently-firing neurons capture much of the changes to and location dependence of population activity in medial prefrontal cortex. This implies that independent neurons can account for much of the population activity structure within each epoch. We take a closer look at this conclusion here.

A useful measure of the overall structure of the population spiking activity is the proportion of “1’s” that encode two or more spikes. The occurrence rates of these “binary errors” across different bin sizes tell us about the burst structure of the neural activity. Figure 8A shows that increasing the bin size applied to the data interpolates between words of single spikes and words of spike bursts in both training and sleep epochs. At bin sizes less than 10 ms, almost all 1’s in each word are single spikes; at bin sizes above 50 ms, the majority of 1’s in each word are two or more spikes and so encode a burst of spikes from a neuron.

**Figure 8.**
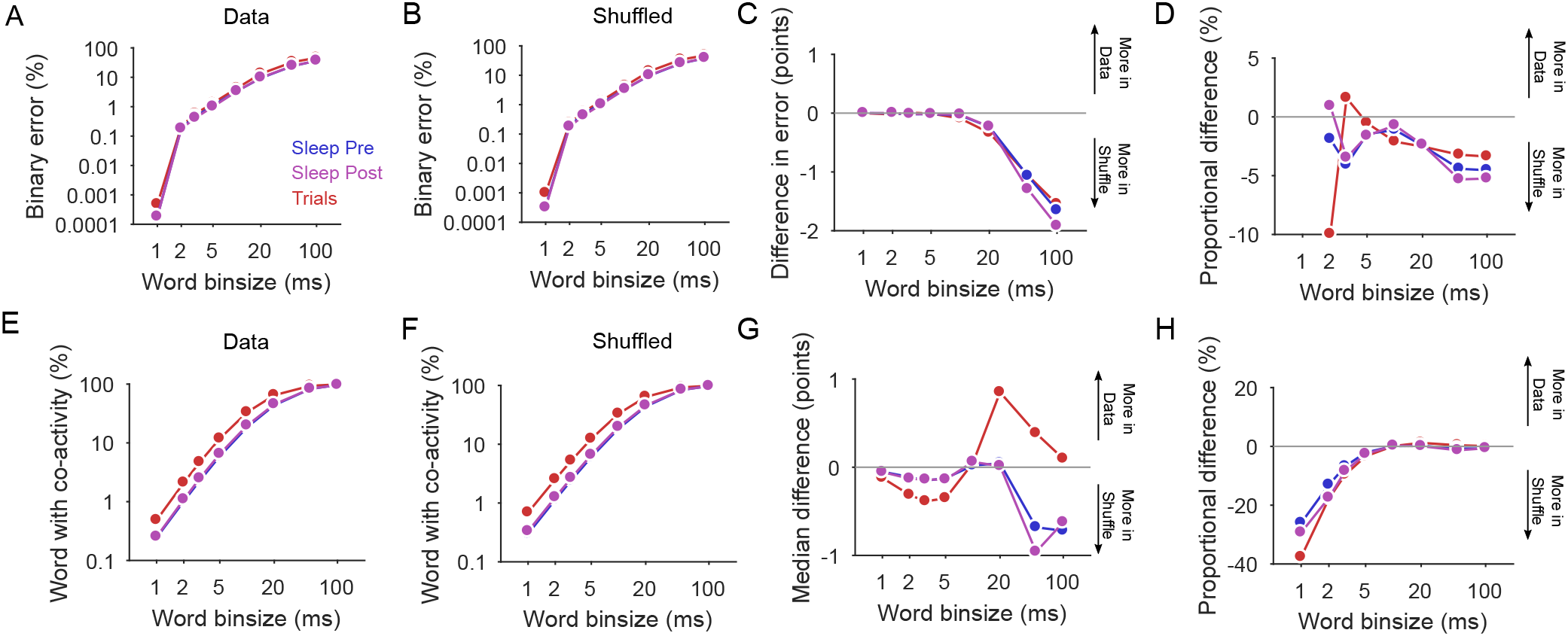
Independent neurons capture a large fraction of population activity structure. (A) Proportion of “1’s” that encode more than one spike (“binary error”), across all emitted words in all learning sessions. Epoch colours apply to all panels. (B) As panel (A), for dictionaries of independent neurons derived by shuffling neuron inter-spike intervals to remove correlations. Proportions are means from 20 shuffled datasets of the learning sessions. (C) Mean difference between binary error proportions in the data and predicted by independent neurons, in percentile points. (D) As panel (C), expressed as a proportion of the binary errors in the data. (E) Proportion of emitted words in each epoch that have more than one active neuron, pooled across all learning sessions (replotted from Figure 5H). (F) As panel (E), for dictionaries of independent neurons. (G) Median difference between the proportion of emitted co-activity words in the data and predicted by independent neurons. (H) As panel (G), expressed as a proportion of the number of co-activity words in the data.

Dictionaries of independent neurons largely recapitulate these bin size dependencies for all epochs (Figure 8B-D). Their only departure is about 5% more binary errors than in the data at bin sizes above 20 ms (Figure 8D). As by construction there are the same number of spikes for each neuron in the data and independent neuron dictionaries, this implies that the data contain more spikes per burst on 50-100 ms time-scales (so that there are fewer bins with bursts in total).

A useful summary of the joint structure of population activity is the fraction of emitted words that code for two or more active neurons. For the data, increasing the bin size increases the fraction of emitted words that contain more than one active neuron (Figure 8E), from about 1% of words at 2 ms bins to all words at 50 ms bins and above. There are consistently more of these co-activity words in training epochs than sleep epochs for the same bin size, pointing to more short time-scale synchronous activity during movement along the maze than in sleep.

Dictionaries of independent neurons also recapitulate these bin size and epoch dependencies of neural co-activity (Figure 8F-H). Figure 8H shows that the independent neuron dictionaries have more co-activity words at small bin sizes. It might be tempting here to conclude that the data dictionaries are constrained to fewer co-activity words than predicted by independent neurons; but these differences are equally consistent with a shadowing effect from spike-sorting, where one or more near-simultaneous spikes from neurons on the same electrode are missed (Harris et al., 2000; Bar-Gad et al., 2001): when the data are shuffled, more near-simultaneous spikes between neurons are possible. Nonetheless, above bins of 5 ms, the disagreement between the data and independent neuron dictionaries is proportionally negligible (Figure 8H). Consequently, much of the population activity in medial prefrontal cortex is well-captured by an independent-neuron model, perhaps pointing to a high-dimensional basis for neural coding.

## Discussion

We studied here how the structure of population activity in medial prefrontal cortex changes during rule-learning. We found the structure of instantaneous population activity in sleep always changes after training, irrespective of any change in overt behaviour during training. This plasticity of population activity could be entirely accounted for by independent changes to the excitability of individual neurons. Unique to learning is that changes to the structure of instantaneous population activity during training are carried forward into the following bouts of sleep. Population plasticity during learning includes both changes to individual neuron excitability and to co-variations of firing rates between neurons. These results suggest two forms of population plasticity in medial prefrontal cortex, one a constant form unrelated to learning, and the other correlated with the successful learning of action-outcome associations.

To isolate learning and non-learning changes, we found useful the “strong inference” approach of designing analyses to decide between simultaneous hypotheses for the same data. We identified separable sessions of learning and stable behaviour in order to contrast the hypothesis that population structure would only change during overt learning against the hypothesis that population structure is always changing irrespective of behaviour. Similarly, we contrasted three hypotheses for what drove those changes in population structure: changes to excitability of independent neurons; changes in brief co-variations of rates; and changes in precise co-spiking.

### A dictionary of cortical activity states

Characterising the joint activity of cortical neurons is a step towards understanding how the cortex represents coding and computation (deCharms and Zador, 2000; Wohrer et al., 2013; Yuste, 2015). One clue is that the joint activity of a cortical population seems constrained to visit only a sub-set of all the possible states it could reach (Tsodyks et al., 1999; Luczak et al., 2009; Sadtler et al., 2014; Jazayeri and Afraz, 2017), in part determined by the connections into and within the network of cortical neurons (Galan, 2008; Marre et al., 2009; Ringach, 2009; Buesing et al., 2011; Habenschuss et al., 2013; Kappel et al., 2015). This view predicts that changing the network connections through learning would change the set of activity states (Battaglia et al., 2005).

We see hints of this prediction in our data. We found changes to the probability of words in training that are detectable in post-training sleep, consistent with the idea that reinforcement-related plasticity of the cortical network has persistently changed the constrained set of activity states. But changing the network’s connections should change not just the set of activity states, but also their sequences or clustering in time (Tkacik et al., 2014; Ganmor et al., 2015). This suggests that further insights into population plasticity with these data could be found by characterising the preservation of word sequences or clusters in time between training and sleep epochs, and comparing those to suitable alternative hypotheses for temporal structure.

### Excitability drives constant population plasticity

A change in the statistics of a population’s neural activity is not in itself evidence of learning (Okun et al., 2012). Indeed, we saw here a constant shifting in statistical structure between sleep epochs, regardless of whether the rats showed any evidence of learning in the interim training epoch. As these shifts between sleep could be seen at all time-scales of words we looked at, and were recapitulated by dictionaries of independent neurons, they are most consistent with a model of independent changes to the excitability of individual neurons.

Excitability changes could arise from the spontaneous remodelling of synaptic connections onto a neuron, whether from remodelling of dendritic spines (Fu et al., 2012; Hayashi-Takagi et al., 2015), or changes of receptor and protein expression within a synapse (Wolff et al., 1995; Ziv and Brenner, 2017). Alternatively, these changes could arise from long-lasting effects on neuron excitability of neuromodulators accumulated in medial prefrontal cortex during training (Seamans and Yang, 2004; Tierney et al., 2008; Dembrow et al., 2010; Benchenane et al., 2011). A more detailed picture of this constant population plasticity will emerge from stable long-term population recordings at millisecond resolution (Jun et al., 2017) of the same prefrontal cortex neurons throughout rule-learning.

### Learning correlates with directional population plasticity

Unique to learning a new rule in the Y-maze was that changes to word probability in training were carried forward to post-training sleep. As this persistence of word probability occurred most clearly for short time-scale words (20 ms or less), and were partly driven by changes in rate co-variations, it is most consistent with a model of synaptic changes to the prefrontal cortex driven by reinforcement. A possible mechanism here is that reinforcement-elicited bursts of dopamine permitted changes of synaptic weights into and between neurons whose co-activity preceded reward (Izhikevich, 2007; Benchenane et al., 2011). Such changes in synaptic weights would also alter the excitability of the neuron itself, accounting for the changes between pre and post-training sleep epochs in learning sessions.

A particularly intriguing question is how the constant and learning-specific plasticity of population activity are related. Again, stable long-term recordings of spiking activity in the same population of neurons across learning would allow us to test whether neurons undergoing constant changes in excitability are also those recruited during learning (Lee et al., 2012; Hayashi-Takagi et al., 2015). Another question is how the carrying forward of training changes of population activity into sleep depends on an animal’s rate of learning. In each learning session here the identified learning trial was before the half-way mark, meaning that the majority of words contributing to the training dictionary came from trials after the rule was acquired. It is an open question as to whether the same relationship would be seen in sessions of late learning, or in tasks with continual improvement in performance rather than the step changes seen here.

### Replay and dictionary sampling

The increased similarity of word probability in training and post-training sleep suggests an alternative interpretation of “replay” phenomena in prefrontal cortex (Euston et al., 2007; Peyrache et al., 2009). Replay of neural activity during waking in a subsequent episode of sleep has been inferred by searching for matches of patterns of awake activity in sleep activity, albeit at much coarser time-scales than used here. The better match of waking activity with subsequent sleep than preceding sleep is taken as evidence that replay is encoding recent experience, perhaps to enable memory consolidation. However, our observation that the probabilities of words in stable sessions’ trials are not systematically closer to those in post-training sleep (Figure 4) is incompatible with the simple replay of experience-related activity in sleep. Rather, our results suggest learning correlates with persistent changes to the cortical network, such that words have more similar probabilities of appearing in training and post-training sleep than in training and pre-training sleep. In this view, replay is a signature of activity states that appeared in training being resampled in post-training sleep (Battaglia et al., 2005).

### Population coding of statistical models

What constraints do these changes to mPfC population activity place on theories for acquiring and representing statistical models of actions and their outcomes? In this view, the joint activity of the population during the trials represents something like the joint probability *P*(*a, o|state*) of action *a* and outcome *o* given the current state of the world (Alexander and Brown, 2011); or, perhaps more generally, a model for the transitions in the world caused by actions, *P(stαte(t + 1)|α, state(t))*. Such models could support the proposed roles of medial prefrontal cortex in guiding action selection (by querying the outcomes predicted by the model), or monitoring behaviour (by detecting unexpected deviations from the model). The changes in the structure of population activity during learning are consistent with updating such models based on reinforcement.

Our results show these dictionary changes are carried forward to the spontaneous activity of sleep, suggesting that the encoded statistical model is present there too. One explanation for this stems from the sampling hypothesis for probability encoding. In this hypothesis, a population encodes a statistical model in the joint firing rates of its neurons, so that the pattern of activity across the population at each moment in time is a sample from the encoded distribution (Fiser et al., 2010; Berkes et al., 2011). This hypothesis predicts that spontaneous activity of the same neurons must still represent samples from the statistical model: but in the absence of external input, these are then samples from the “prior” probability distribution over the expected properties of the world.

According to this hypothesis, our finding that learning-driven changes to population structure are conserved in post-training sleep is consistent with the statistical model now reflecting well-learnt expected properties of the world – namely, that a particular set of actions on the maze reliably leads to reward. In other words, the prior distribution for the expected properties of the world has been updated. Further, the sampling hypothesis also proposes a role for the constant changes of excitability without obvious direction – that such spontaneous plasticity explores possible configurations of the network and so acts as a search algorithm to optimise the encoded statistical model (Kappel et al., 2015; Maass, 2016). These links, while tentative, suggest the utility of exploring models for probabilistic codes outside of early sensory systems (Fiser et al., 2010; Pouget et al., 2013).

## Acknowledgments

We thank Silvia Maggi and Rasmus Petersen for comments on early drafts of this manuscript, and the Humphries lab (Javier Caballero, Mat Evans) for discussions. A.S. and M.D.H were supported by a Medical Research Council Senior non-Clinical Fellowship award MR/J008648/1 to M.D.H, and Medical Research Council Grant MR/P005659/1. A.P. was supported by a Canada Research Chair Tier 2 (154808). The original data were obtained through funding from the EU Framework 6 “ICEA” project.

